# Nanopore Sequencing at Mars, Europa and Microgravity Conditions

**DOI:** 10.1101/2020.01.09.899716

**Authors:** Christopher E. Carr, Noelle C. Bryan, Kendall N. Saboda, Srinivasa A. Bhattaru, Gary Ruvkun, Maria T. Zuber

## Abstract

Nanopore sequencing, as represented by Oxford Nanopore Technologies’ MinION, is a promising technology for *in situ* life detection and for microbial monitoring including in support of human space exploration, due to its small size, low mass (∼100 g) and low power (∼1W). Now ubiquitous on Earth and previously demonstrated on the International Space Station (ISS), nanopore sequencing involves translocation of DNA through a biological nanopore on timescales of milliseconds per base. Nanopore sequencing is now being done in both controlled lab settings as well as in diverse environments that include ground, air and space vehicles. Future space missions may also utilize nanopore sequencing in reduced gravity environments, such as in the search for life on Mars (Earth-relative gravito-inertial acceleration (GIA) *g* = 0.378), or at icy moons such as Europa (*g* = 0.134) or Enceladus (*g* = 0.012). We confirm the ability to sequence at Mars as well as near Europa or Lunar (*g* = 0.166) and lower *g* levels, demonstrate the functionality of updated chemistry and sequencing protocols under parabolic flight, and reveal consistent performance across *g* level, during dynamic accelerations, and despite vibrations with significant power at translocation-relevant frequencies. Our work strengthens the use case for nanopore sequencing in dynamic environments on Earth and in space, including as part of the search for nucleic-acid based life beyond Earth.

## Introduction

Life as we know it uses deoxyribonucleic acids (DNA) as the basis for heritability and evolution. Life beyond Earth might utilize identical or similar informational polymers due to the widespread synthesis of common building blocks, common physicochemical scenarios for life’s origin(s), or common ancestry via meteoritic exchange, most plausible for Earth and Mars. Beyond the search for life, sequencing is of high relevance for supporting human health on Earth and in space, from detecting infectious diseases, to monitoring of biologically-based life support systems.

Nanopore sequencing^1^, as commercialized by Oxford Nanopore Technologies, is a promising approach that is now used ubiquitously in the lab and in the field. McIntyre et al. (2016) reported a single mapped read obtained via nanopore sequencing during parabolic flight, obtained across multiple parabolas^2^. Vibration of flow cells revealed that 70% of pores should survive launch, consistent with later successful nanopore sequencing on the ISS^3^. However, we are not aware of any nanopore experiments that attempted to quantify the impact of vibration while sequencing.

Here we test the impacts of: 1) altered *g* level, 2) vibration, and 3) updated chemistry/flow cells.

## Results

Flight operations were conducted on November 17, 2017 onboard a Boeing 727-200F aircraft (G-Force One®, Zero Gravity Corporation). Four sets of parabolas were performed with 5, 6, 4, and 5 parabolas respectively (**Fig. 1a**). The first set targeted, in order, Mars *g*, Mars *g*, Lunar *g*, 0 *g*, and 0 *g* (**Fig. 1b**). All other parabolas targeted 0 *g*. The flight profile was segmented into periods of “transition,” “parabola,” “hypergravity,” and “other” (typically, gentle climb, descent, straight and level flight, or standard rate turns) on the basis of accelerometer measurements^4^. Sequencing was also performed on the ground prior to the flight as a control.

**Fig. 1.**
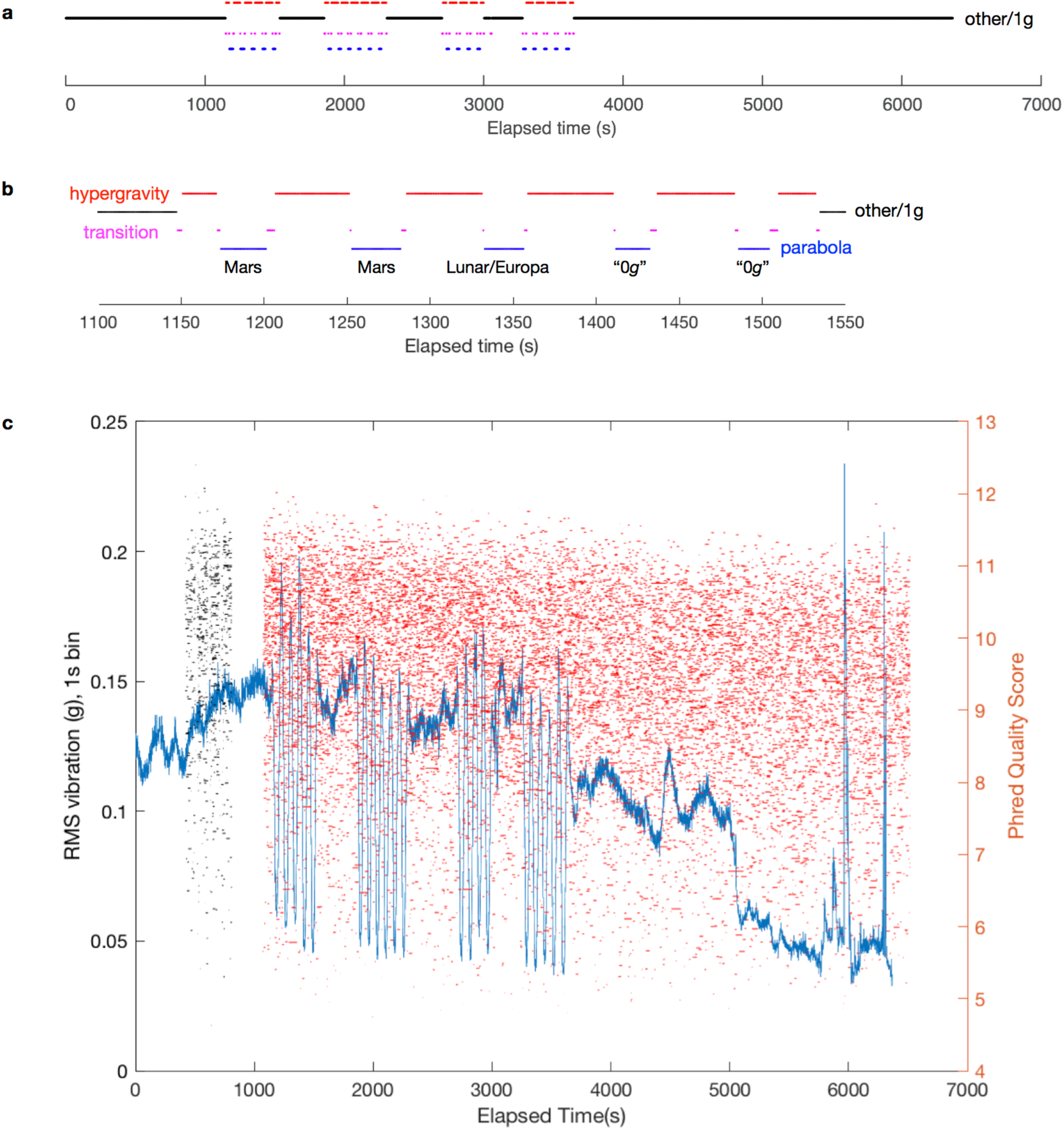
Single molecule sequencing during parabolic flight. **a** Phases of flight timeline (black: other/1g; red: hypergravity; magenta: transition; blue: parabola). **b** Phases of flight for first set of parabolas. **c** Vibration (blue line, left axis) and sequencing reads measured during flight; each read is represented by a horizontal line (mux=black, run=red) at its representative read quality score,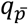.

### Sequencing

Sequencing of control lambda DNA was performed for a total of 38 minutes on the ground and 103 minutes during flight, on the same flow cell, resulting in 5,293 and 18,233 reads for ground (**Supplementary Fig. 7**) and flight (**Fig. 1c; Supplementary Fig. 7**) respectively, of which 5,257 and 18,188 were basecalled (**Supplementary Tables 1-2**). Of the flight reads, 14,431 fell wholly within a phase of flight, including parabola (404), hypergravity (1996), transition (7), and other (12,024). Sequencing reads were obtained during all parabolas, including under Mars, lunar/Europa, and zero-*g* conditions (**Fig. 2**). The *g* levels achieved during each parabola were previously reported^4^. For the purposes of statistical analysis, mux reads (**Fig. 1c**, black horizontal lines) were excluded to avoid any sequencer start-up effects.

**Fig. 2.**
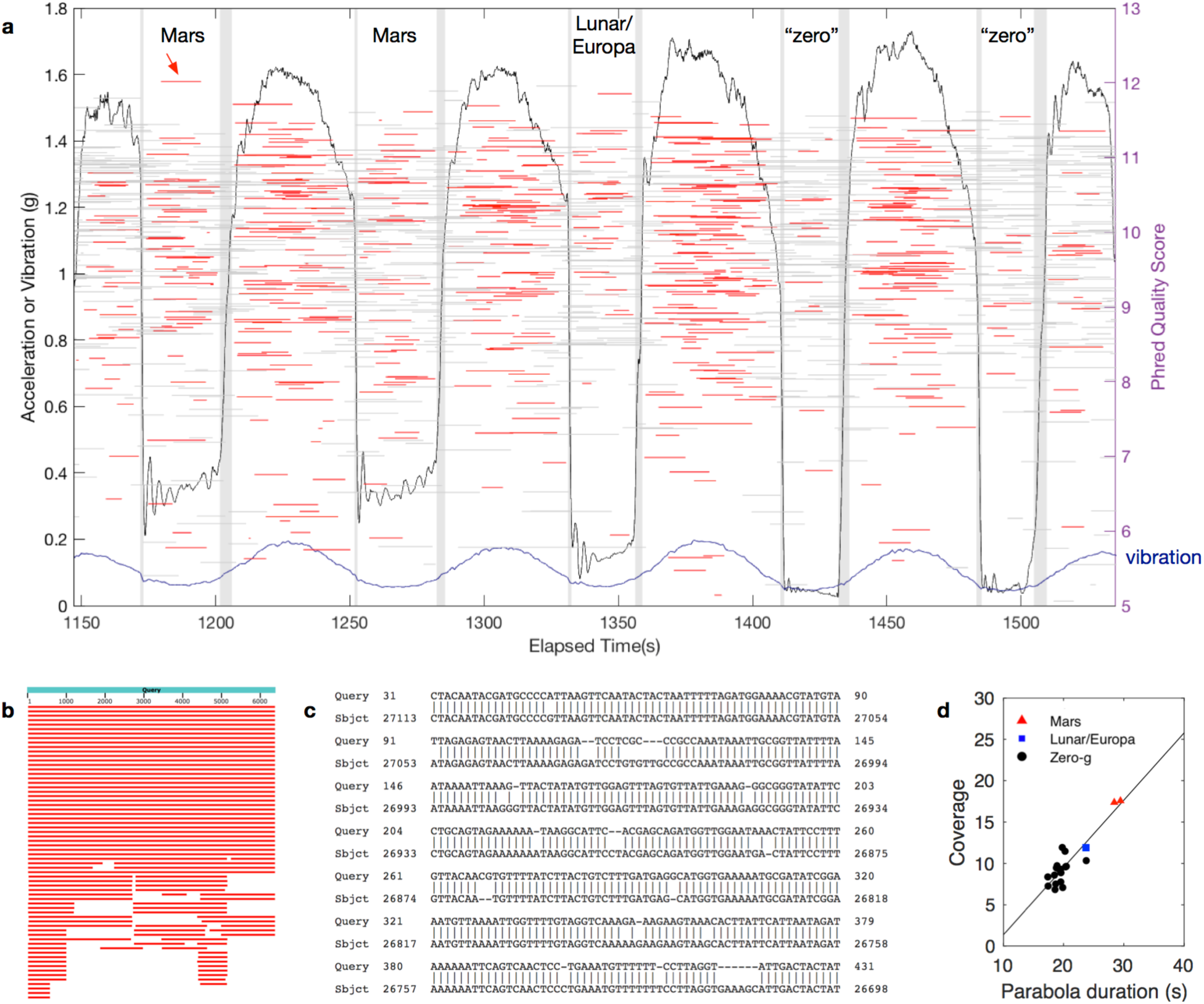
Sequencing in reduced gravity. **a** *g* level achieved (black line) and RMS vibration (1 s bins, blue line) and associated sequencing reads acquired during first “Mars” parabola. Each read is represented by a horizontal line (grey: partially or completely in transition period; red: completely in non-transition period) at its representative read quality score, 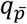 (right axis). Vertical gray bands demarcate transitions between phases of flight. **b** Top scoring BLAST results for highest quality “Mars” read, indicated via arrow in panel *a*, length 6402. **c** Start of best match sequence alignment, to J02459.1 *Enterobacteria phage lambda*, complete genome, length 48502 (range 20562 to 27113, score 8907 bits(9877), expect 0.0, identities 6108/6651 (92%), gaps 395/6651 (5%), strand Plus/Minus). **d** Average genomic coverage of lambda for all parabolas based on tombo-aligned bases.

### Vibration

Zero-phase filtering effectively removed frequencies at or below 10 Hz (**Supplementary Fig. 2-3**). Filtered root-mean-square (RMS) vibration varied throughout the flight and showed clear deviations associated with parabolas (**Fig. 1c**; **Fig. 2a**), indicating a smoother environment during freefall. Remaining aircraft-associated vibrations were largely in the 10 Hz to 1 kHz band with peaks at 116-128 Hz, 250-270 Hz, 495-496 Hz, 580-680 Hz, 876 Hz (**Supplementary Fig. 3**). During zero-*g* parabolas, the magnitude of the residual *g* level and vibrations were comparable (**Fig. 2a**).

### Integrated Read-Level Analysis

Stepwise linear regression was used to determine whether time and RMS vibration could predict median sequence quality (**Supplementary Fig. 7**), the Phred quality score^5,6^ associated with the average per base error probability of a given read (See Materials and Methods). Unlike ground operations, where time was the only significant predictor of sequence read quality (*p* = 0), time, *g* level, and their combined effects were predicted to be significant indicators during flight (all *p* =< 10^−4^; **Supplementary Tables 3-4**). However, in both cases, the variance explained was small (*adj. R*^2^ = 0.060 and 0.275, respectively, for ground and flight).

In order to elucidate the role of *g* level on read quality, those reads falling wholly within an individual phase of flight were examined using a one-way ANOVA, with Tukey’s Honest Significant Difference (HSD) post-hoc analyses (**Supplementary Table 5**). Sequence quality was significantly different during each phase of flight, with the lowest read quality during parabolas 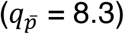 and the highest quality 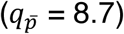 during hypergravity (**Supplementary Fig 10**).

### Integrated Base-Level Analysis

Tombo^7^ was used to associate raw ionic current signals with specific genomic bases, and the number of reads aligning was similar to the number of reads with Phred quality scores^5,6^ > 6.5. The percentage of bases that aligned to the lambda genome via tombo^7^ was 87.8% and 89.7% for ground and flight, respectively (**Supplementary Table 1**). Average coverage for tombo-aligned bases was adequate to sequence the lambda genome many times over during each parabola (**Fig. 2d**) and the coverage was largely explained by parabola duration (*adj. R*^2^ = 0.807; **Supplementary Table 9**).

By aligning ionic current signals to bases, tombo allowed us to measure the translocation time associated with each base (**Supplementary Fig. 6**), the time required for the motor protein, acting as a rachet, to move the DNA strand one base into the nanopore. Translocation here refers to motion of the motor protein relative to the DNA strand, and not the total time to get through the nanopore, which requires many translocation steps. The inverse of translocation time is a direct measurement of sequencing rate for a given nanopore (bases/s).

Despite the nearly 6-fold (5.89) average higher RMS vibration during flight compared to ground (**Fig. 3a**), the probability densities for translocation time are strikingly similar (**Fig. 3b; Supplementary Fig. 6**). However, base translocation times were significantly different (Kolmogorov-Smirnov, two-tailed, *p* = 0, test statistic 0.0306), with a slight shift towards longer translocation times during flight. Notably, the median base translocation times were identical (7 samples or 1.8 ms) and the means only differed by 0.125 ms (2.2786 ms, ground; 2.4035 ms, flight). Thus, translocation times were robust to large variations in vibration.

**Fig. 3.**
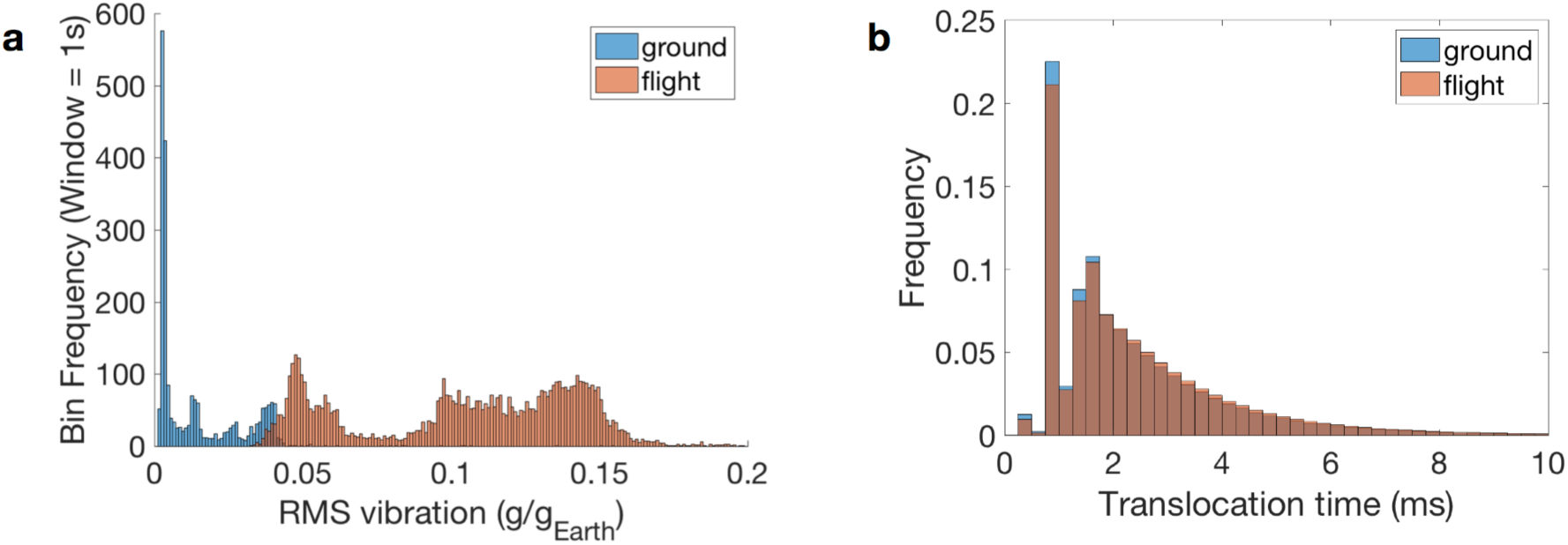
Translocation time is weakly or not affected by vibration. **a** RMS vibration distributions for ground and flight. **b** Nanopore translocation time as measured by alignment of ionic current to the genomic reference: distribution for <10 ms. Ground (blue), flight (light brown), both (dark brown).

Ionic current noise is the variation in the flow of ions passing through the nanopore, measured here at the per-base level as a normalized signal standard deviation determined by tombo through optimal alignment of measured ionic current to a genomic sequence^7^. A stepwise linear regression was performed to determine if time, RMS vibration, or their combined effects were significant predictors of ionic current noise during ground (**Supplementary Fig. 8; Supplementary Table 6**) and flight (**Supplementary Fig. 8; Supplementary Table 7**) operations. Flight analysis included the additional variable *g* level.

For ground operations, the impact of time alone was not significant. However, both vibration (*p* = 0.0018) and the interaction effect of time and vibration (*p* = 0.041) were significant predictors of ionic current noise (**Supplementary Table 6**). However, the explanatory power of the regression was low (*adj. R*^2^ = 0.009). Conversely, time was the only significant predictor of the effect on ionic current noise during flight. Neither RMS vibration, *g* level, nor any of their respective combined effects had significant impacts on ionic current noise (**Fig. 4**; **Supplementary Table 7**).

**Fig. 4.**
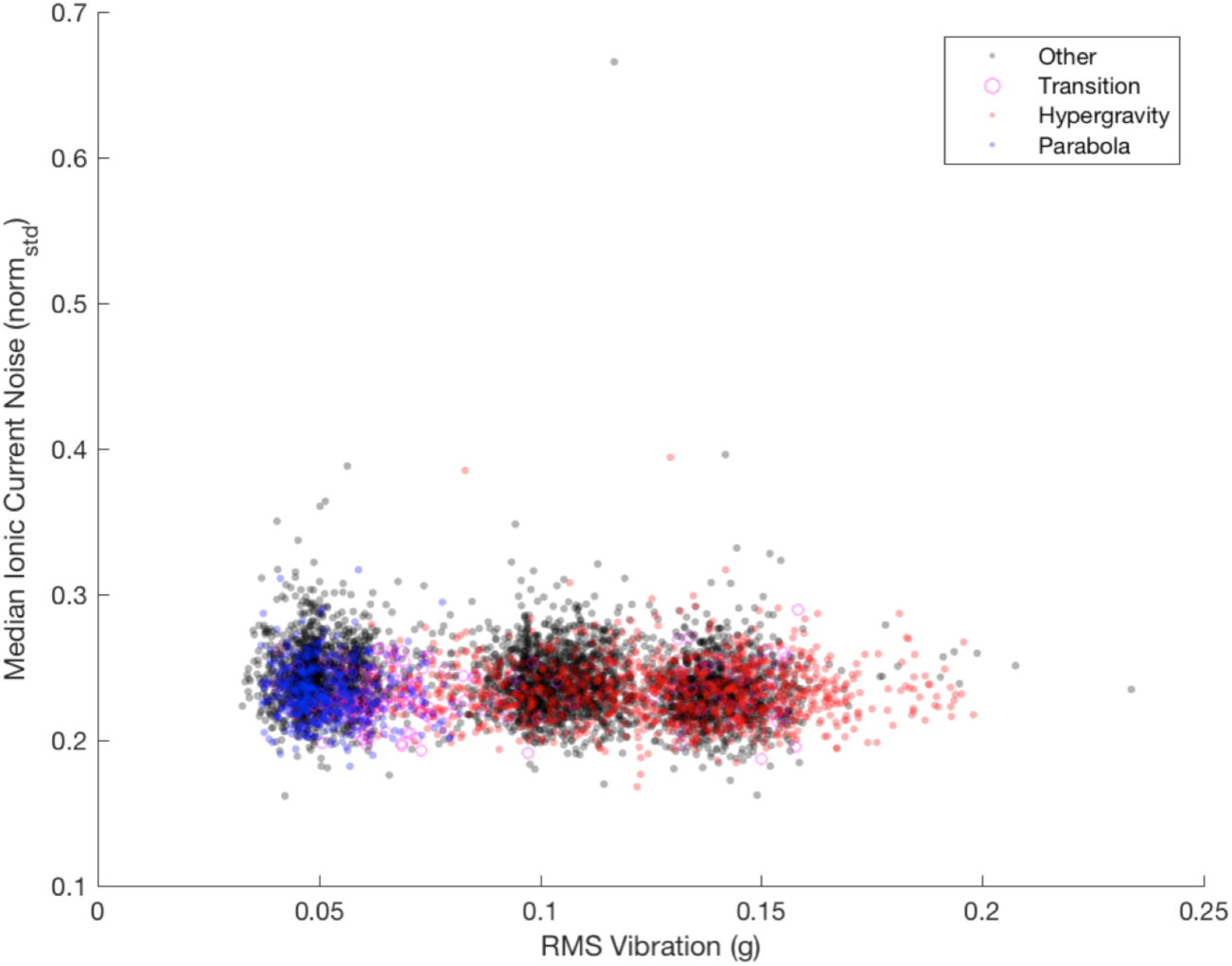
RMS vibration and Median Ionic Current Noise During Flight. The single 1 s period with median ionic current noise > 0.5 has a median absolute deviation (MAD) of > 15 and is therefore an outlier (typically defined as MAD>3).

Because time was a significant indicator of ionic current noise during flight, it was necessary to assess whether the effect could be attributed to a specific phase of flight (**Supplementary Table 8**). Tukey’s HSD post-hoc test demonstrated that out of all six possible pairwise comparisons, only one, parabola vs. transition, was not significant (*p* = 0.345; other *p* < 10^−3^). Ionic current was significantly lower in hypergravity, parabola, and transition phases as compared to other. Ionic currents during hypergravity phases were, on average, lower than all other phases (**Supplementary Table 8**; **Supplementary Fig. 10**). Thus, while the impact of phase of flight on read quality showed a trend towards higher read quality with higher *g* level (**Supplementary Fig. 9**), no such pattern was observed with ionic current (**Supplementary Fig. 10**).

## Discussion

The Mars 2020 rover, currently in development, is expected to touch down in Jezero Crater in February, 2021. While this mission will not attempt to detect extant life, it represents a new era in the search for life beyond Earth. Ambiguous or positive results in the search for ancient life could usher in a new era of life detection efforts, including instrumentation aimed at measuring the presence of nucleic acids, one of the “smoking gun” pieces of evidence for life beyond Earth^8^. In preparation for future life detection missions targeting DNA, we explored the capabilities of nanopore sequencing, and present results demonstrating its successful performance while experiencing aircraft vibrations and under altered *g* levels, including those that would be encountered on the surface of Mars, the Moon, and/or Jupiter’s moon Europa. Due to the limitations of parabolic flight, our zero-*g* conditions involved mean acceleration around 4X higher (0.041 ± 0.005 g)^4^ than at the surface of Saturn’s moon Encealdus (0.011 *g*).

Several factors may have influenced the overall quality of our nanopore sequencing data. The DNA sequencing library used in this experiment was stored for 72 h prior to loading onto the flow cell. As such, the sample was potentially subjected to degradation, which could impact read quality, and may have resulted in the loss of ligated adaptors. Such conditions would negatively impact the proper loading of DNA into the individual nanopore. In addition, there is an expected degradation of the flow cell over time during sequencing, which could explain some of the time-related trends, independent of any effects of vibration or acceleration. Despite sequencing for a limited time at any given *g* level during parabolic flight, the operation of the MinION for sustained periods on the ISS^3^ gives us confidence that extended periods of reduced *g* level does not negatively impact nanopore sequencing. In addition, it provides confidence in nanopore sequencing as a viable life detection technology in very low but non-zero *g* environments, such as Enceladus.

Because zero-phase filtering of vibration data effectively removed frequencies at or below 10 Hz (**Supplementary Fig. 2-3**), filtered vibration measurements did not reflect frequencies where sensor data would be inaccurate due to the non-unity frequency response of the sensor near DC (0 Hz). In addition, this filtering ensured that we could assess the independent effects of *g* level and vibration.

The peaks in the vibration spectrum occurs at frequencies relevant to nanopore sequencing. Despite this, vibration did have any significant impacts on sequence quality nor on ionic current noise, except during ground-based sequencing, where the explanatory power of vibration was negligible (< 1%; **Supplementary Table 6**).

Random vibration at translocation-relative frequencies could exert a minor interfering effect on translocation, although any impact in changes in vibration during flight did not translate into any consistent or large changes in ionic current noise due to the small (0.125 ms) mean difference in translocation times despite nearly a 6-fold change in RMS vibration.

Higher *g* levels tended to be associated with higher read quality, although the effect size is small (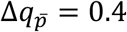, hypergravity – parabola; **Supplementary Table 8**, an upper bound of ∼0.2/*g*). The smallest mean values of ionic current noise were also observed during hypergravity (0.281; **Supplementary Fig. 10).** Although statistically larger, the difference between largest mean value for ionic current noise (phase other) was miniscule (0.003).

Nanopore sequencing is compatible with many life detection missions from the perspective of mass (∼100 g), size, and power (<2 W). Recent work also suggests that MinION electronics and flowcell components would survive radiation doses consistent with life detection missions to Mars, Venus, and Enceladus, although not Europa, without additional shielding^9^. Our work shows that sequencing on all these worlds, including Europa, could be feasible from a *g* level perspective. In addition, the robustness to vibration suggests that operation concurrent with other mission activities, such as drilling or operation of other instrument payloads, could occur without any substantial negative impacts. In addition, our work highlights the potential for nanopore sequencing on Earth and beyond in mobile and dynamic environments such as on passenger aircraft, drones, wheeled vehicles, ships, buoys, underwater vehicles, or other platforms.

## Methods

### Acceleration Measurement and Flight Profile Segmentation

The flight profile was segmented as described in Carr et al.^4^ from acceleration data collected using a metal-body Slam Stick X™ (Mide Technology Corp.). The accelerometer was mounted next to the MinION to a common baseplate, using double sided sticky tape (3M 950) to provide a near-unity vibration frequency response. Vibrations were measured with the internal triaxial piezoelectric accelerometer (TE Connectivity Ltd., 832M1) at a frequency of 5 kHz.

### Sequencing

Sequencing libraries were prepared using DNA derived from *Enterobacteria phage lambda* (NEB N3011S), fragmented using a g-TUBE™(Covaris® 520079) with the 6 kb protocol. Next, the libraries were prepared using the 1D ligation method (SQK-LSK108) using a “one-pot” barcoding protocol^10^ and stored at 4°C for ∼72 hours prior to the flight. At the time of storage, the total library DNA was estimated to be 440 ng at 31.4 ng/ul as assessed by fluorometry (ThermoFisher Qubit® 3.0 Fluorometer with Qubit™ dsDNA HS Assay Kit, Q32854).

A flowcell (FLO-MIN106 R9) was loaded on the ground and sequencing performed using an offline version of MinKNOW 1.7.14 in the flight hardware configuration while the aircraft was on the ground. After 38 minutes, sequencing was stopped. In flight, sequencing was reinitiated around 12 minutes prior to parabolic flight maneuvers, and continued for a total of 103 minutes before termination. After the flight, basecalling was performed with ONT Albacore version 2.3.1 with quality filtering disabled.

### Sequence Data Processing

To quantify adaptor sequences, fastq output was trimmed using Porechop (https://github.com/rrwick/Porechop docker container quay.io/biocontainers/porechop:0.2.3_seqan2.1.1--py35_2). Original untrimmed fast5 reads were aligned to the reference genome (NEB Lambda, equivalent to NCBI NC_001416.1 with mutations 37589 C->T, 45352 G->A, 37742 C->T and 43082 G->A) using tombo (docker container quay.io/biocontainers/ont-tombo: 1.5--py27r351h24bf2e0_0)^7^ with the --include-event-stdev option.

### Sequencing and Acceleration Data Integration

A custom script was used to parse tombo-processed fast5 files to characterize each read and each tombo-aligned genomic base within each read (Supplementary Data). A representative read quality score was calculated as 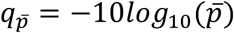, where 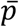 is the mean of the per base error probability *p* = 10^−*q*/10^, where *q* is the per base Phred quality score^5,6^ estimated via basecalling. Read timings were adjusted by offsets to align genomic and accelerometer data (**Supplementary Table 2**). Each read and base was assigned one of the following states (parabola, transition, hypergravity, other) on the basis of the periods.txt file produced by prior analysis^4^ and available online at https://osf.io/nk2w4/.

### Vibration Data Processing

A vibration equivalent to *g* level (Earth relative gravity) was computed as 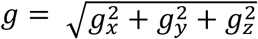 to provide a measure of vibration that is independent of the Slam Stick X™ orientation.

The vibration power spectral density (PSD) for *g* was computed using Welch’s method (MATLAB *pwelch()* function) with default parameters (**Supplementary Fig. 1**). Filtering was then performed for two reasons: 1) to eliminate vibration data where the frequency response of the piezoelectric accelerometer is not unity, and 2) to analyze vibration at frequencies related to timescales at which base translocation occurs during nanopore sequencing, which are overwhelmingly < 10 ms (**Supplementary Fig. 6-7**). The *g* level equivalent vibration *g* was filtered with a high pass infinite impulse response filter (**Supplementary Fig. 2**) that was generated with MATLAB’s *designfilt()* function (stopband 5 Hz @ 60 dB attenuation, passband 10 Hz with unity ripple, sample rate 5kHz). Filtering was performed using the MATLAB *filtfilt()* function, which uses forward and reverse filtering to achieve zero phase delay. The PSD was computed as before for the resulting filtered *g* level equivalent vibration *g*_*f*_ (**Supplementary Fig. 3**). RMS vibration was computed in 1-s bins from *g*_*f*_ using the MATLAB *rms()* fuction. An overview of vibration is shown in **Supplementary Fig. 4** for flight and **Supplementary Fig. 5** for ground.

### Sequencing Read Quality Regression Analysis

Sequencing read times were adjusted by an offset to place sequencing reads into the accelerometer elapsed time (**Supplementary Table 2**). A time series of median read quality was estimated in 1s bins by computing the median of 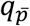 for all reads covering the bin. Stepwise linear regression, via the MATLAB *stepwiselm()* function, was used to evaluate the impact of time, RMS vibration, and *g* level (flight only) on sequence quality, as measured by median 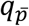 (**Supplementary Tables 3-4**). For flight, the regression time was restricted to a maximum elapsed time of 4000 seconds to eliminate potential confounding effects of the aircraft descent and landing.

### Sequencing Read Quality Phase of Flight Analysis (Flight Only)

To assess differences in read quality as a function of phase of flight, we performed a one-way analysis of variance (ANOVA) via the MATLAB *anova1()* function on the non-mux reads (**Supplementary Table 5**), excluding reads in transition periods due to their low number (7) and short length. To compare group means we then used Tukey’s Honestly Significant Difference test (MATLAB *multcompare()* function), which is conservative for one-way ANOVA with different sample sizes.

### Coverage of Genomic-Aligned Bases

Base times were adjusted by an offset to place each tombo-aligned base into the accelerometer elapsed time (**Supplementary Table 2**). Coverage was estimated as the sum of tombo-aligned bases within a given phase of flight divided by the lambda genome size (48502 bases). Stepwise linear regression, via the MATLAB *stepwiselm()* function, was used to evaluate the relationship between coverage and parabola period.

### Base Ionic Current Noise Regression Analysis

A time series of ionic current noise was estimated in 1s bins by computing the median of ionic current (tombo norm_std output) for all bases within a bin. Stepwise linear regression, via the MATLAB *stepwiselm()* function, was used to evaluate the impact of time, RMS vibration, and *g* level (flight only) on median ionic current noise (**Supplementary Tables 6-7**). For flight, the regression time was restricted as stated above.

### Base Ionic Current Noise Phase of Flight Analysis (Flight Only)

To assess differences in ionic current as a function of phase of flight, we performed a one-way analysis of variance (ANOVA) via the MATLAB *anova1()* function on the non-mux tombo-aligned bases (**Supplementary Table 8**). To compare group means we then used Tukey’s Honestly Significant Difference test as above.

### Does Flight vs. Ground impact Translocation Time?

A Kolmogorov-Smirnov test was performed with the MATLAB *kstest2()* function on the base translocation times for ground vs. flight.

## Code Availability

The MATLAB scripts implementing our analysis are available at: https://github.com/CarrCE/zerogseq

## Data Availability

Raw and calibrated data are available via the Open Science Framework at: https://osf.io/n6krq/ and https://osf.io/nk2w4/

## Acknowledgements

We thank the MIT Media Lab Space Exploration Initiative for providing the parabolic flight.

## Contributions

C.E.C. designed the experiment, C.E.C., K.S., S.A.B. and N.C.B. built and tested the hardware, N.C.B. and M.T.Z. collected the data, C.E.C. and N.C.B. processed the data. G.R. advised on the experiment design. C.E.C. and N.C.B. wrote, and all authors edited and approved, the paper. C.E.C. is the guarantor.

## Supplementary Information

Supplementary information is available at Nature Microgravity’s website.

## Supplementary Information

**Supplementary Fig. 1.**
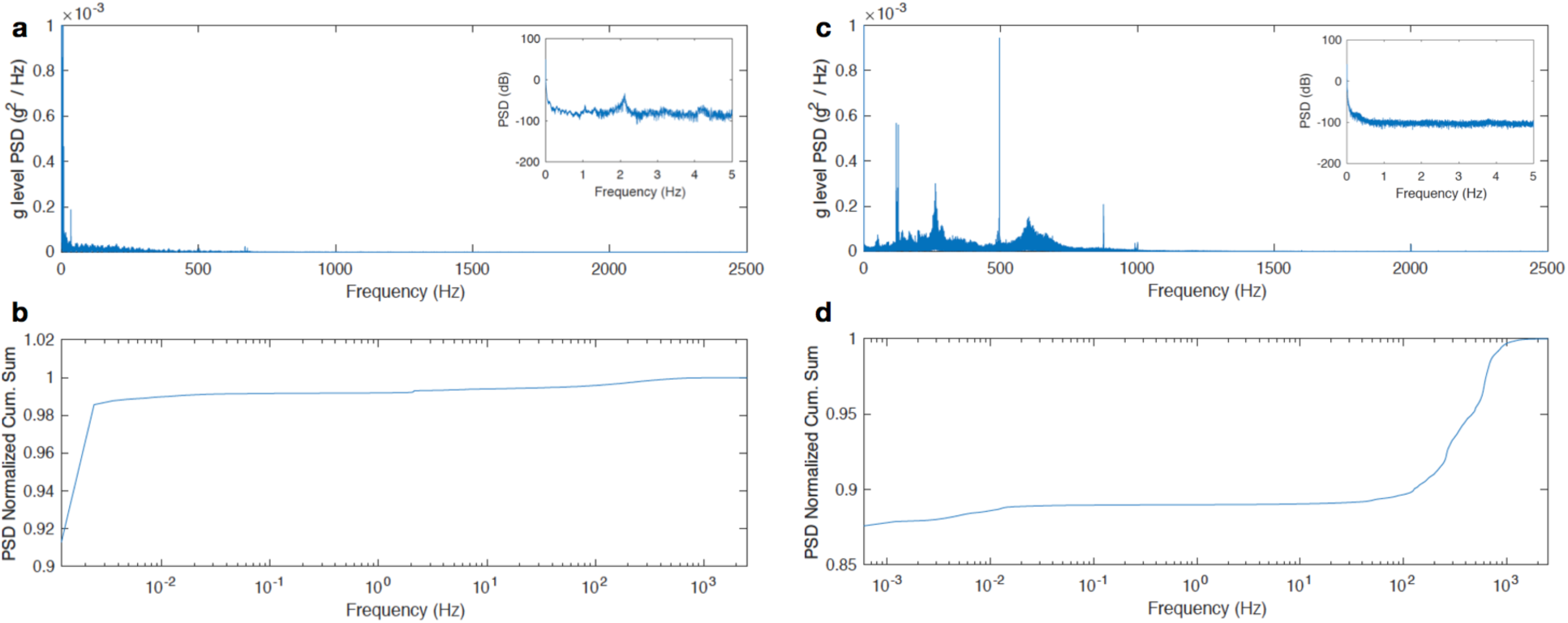
Power spectral density (PSD) of vibration *g*-level equivalent. **a** Ground PSD. **b** Cumulative sum of Ground PSD. **c** Flight PSD. **d** Cumulative sum of Flight PSD.

**Supplementary Fig. 2.**
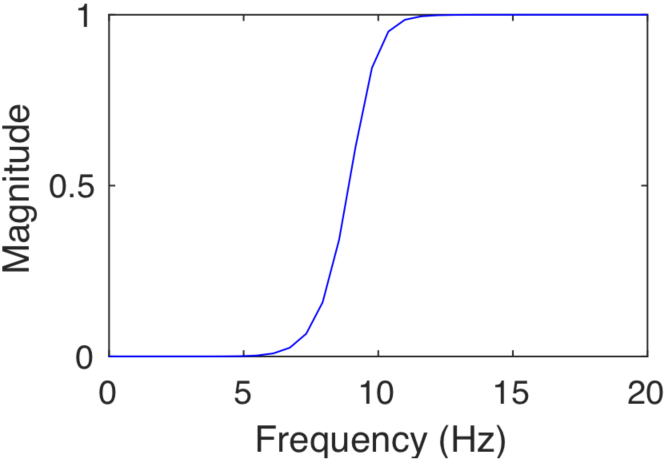
Frequency response of vibration filter. Phase is not shown as it is not relevant due to use of zero-phase filtering.

**Supplementary Fig. 3.**
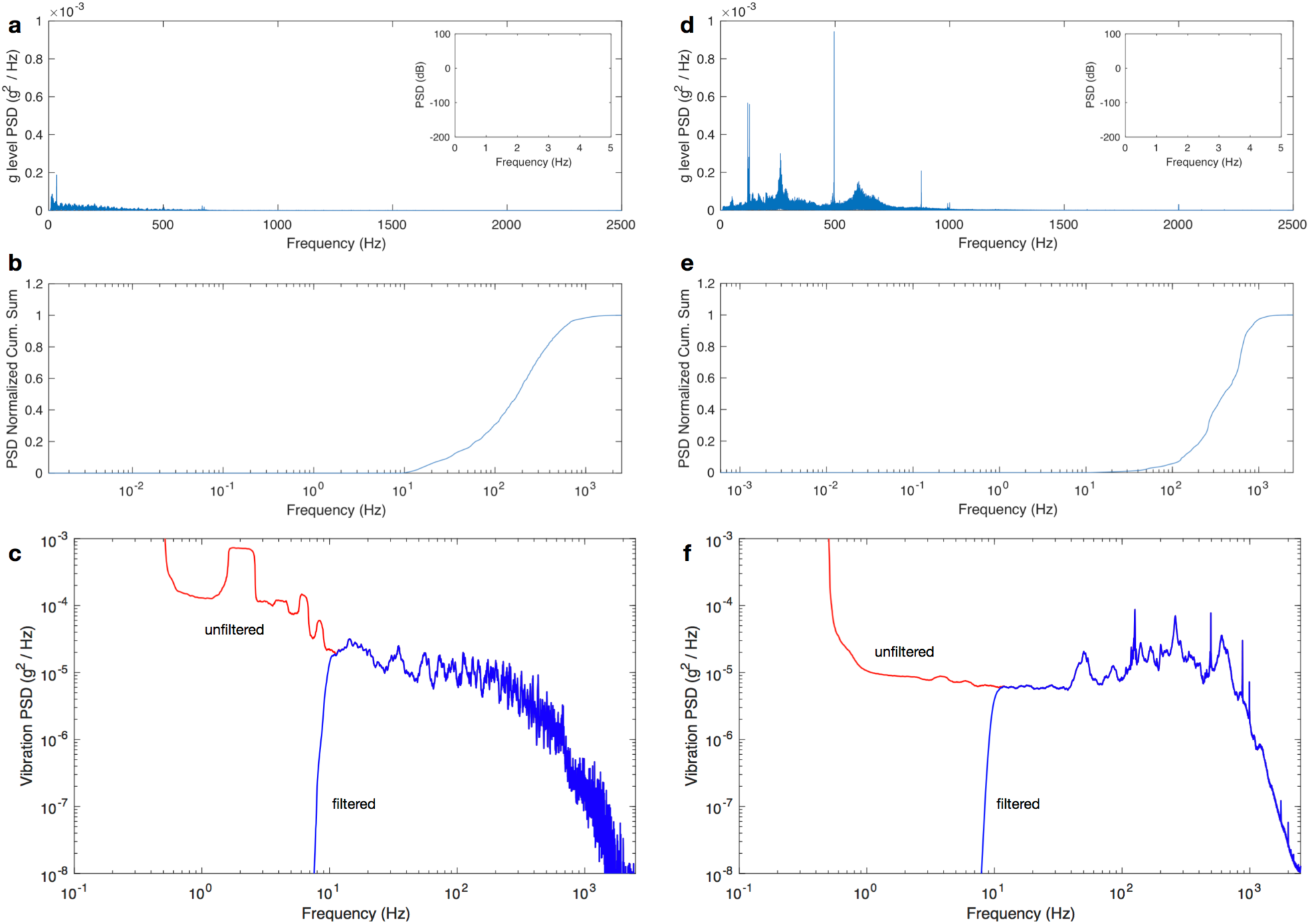
Power spectral density (PSD) of vibration *g*-level equivalent after high-pass filtering. **a** Ground PSD. **b** Cumulative sum of Ground PSD. **c** Ground PSD smoothed with window size of 1 Hz (red=unfiltered, blue=filtered). **d** Flight PSD. **e** Cumulative sum of Flight PSD. **f** Flight PSD smoothed with window size of 1 Hz (red=unfiltered, blue=filtered).

**Supplementary Fig. 4.**
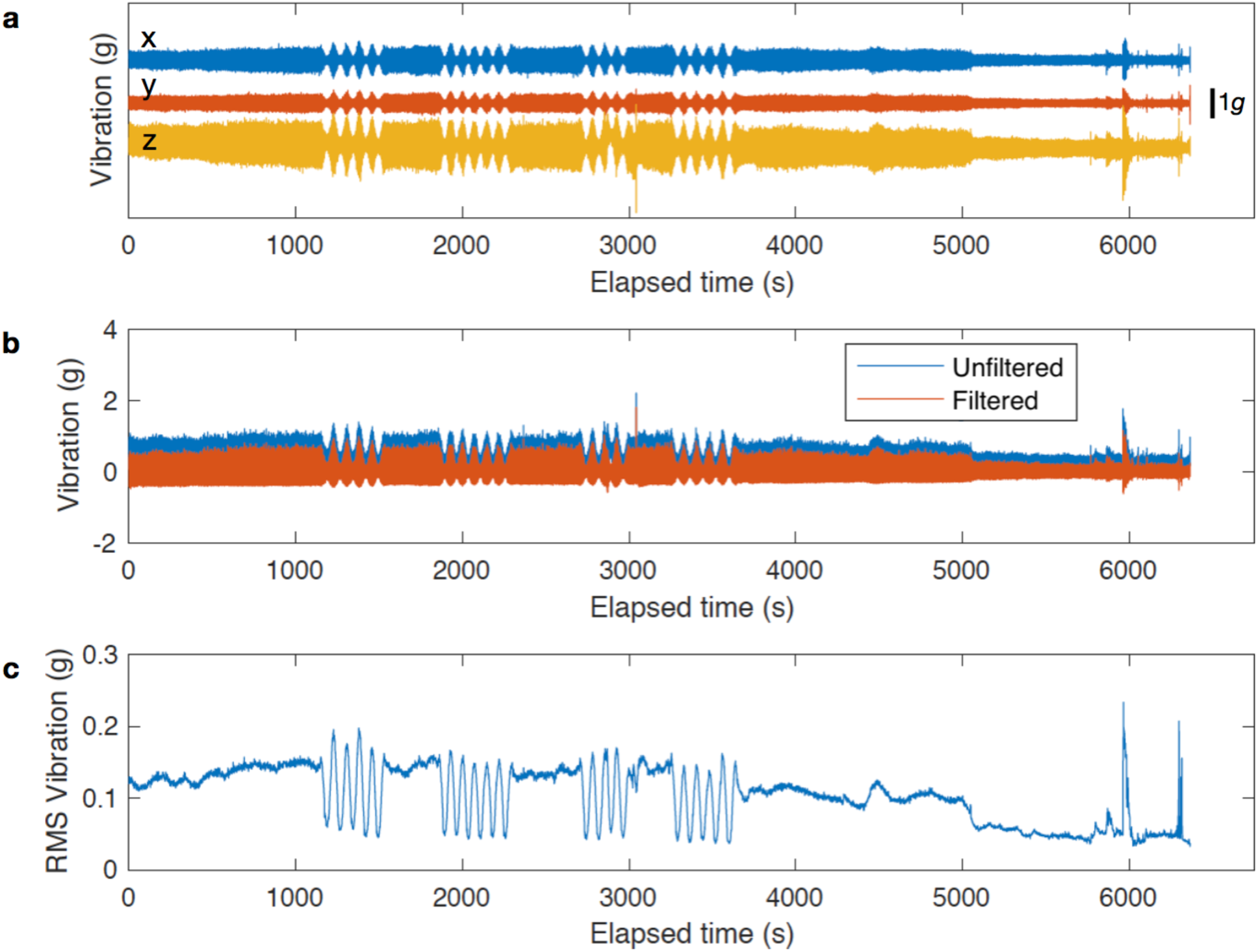
Vibration during flight (overview). **a** Unfiltered vibration measurements after mean removal (1.5 g offsets for display only: x +3 g, y + 1.5 g, z +0 g). Scale bar: 1g. **b** *g*-level equivalent vibration pre- and post-filtering. **c** Root-mean-square (RMS) vibration profile (1 second bin).

**Supplementary Fig. 5.**
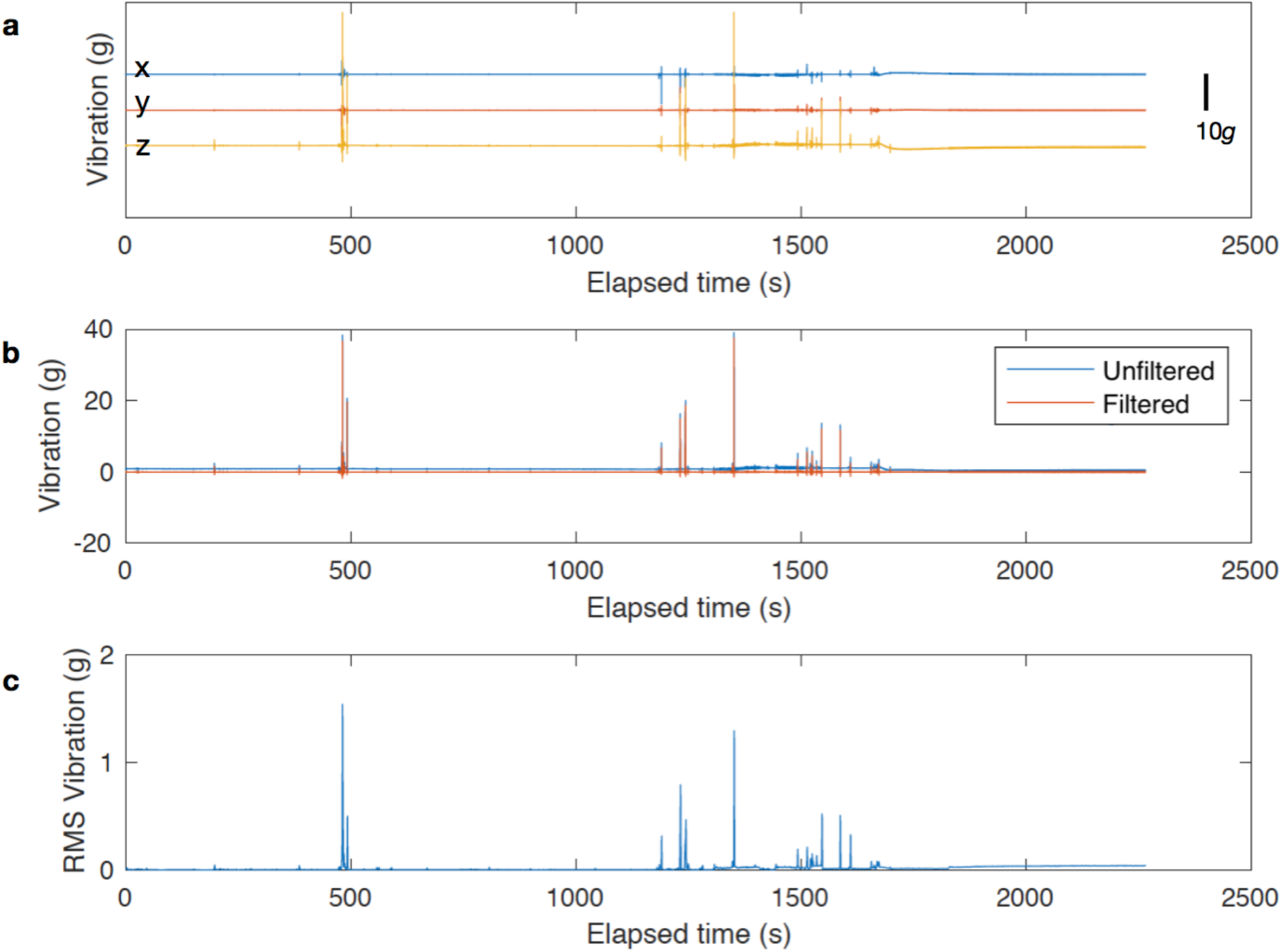
Vibration during ground operations (overview). **a** Unfiltered vibration measurements after mean removal (10g offsets for display). Scale bar: 10g. **b** *g*-level equivalent vibration pre- and post-filtering. **c** Root-mean-square (RMS) vibration profile (1 second bin).

**Supplementary Fig. 6.**
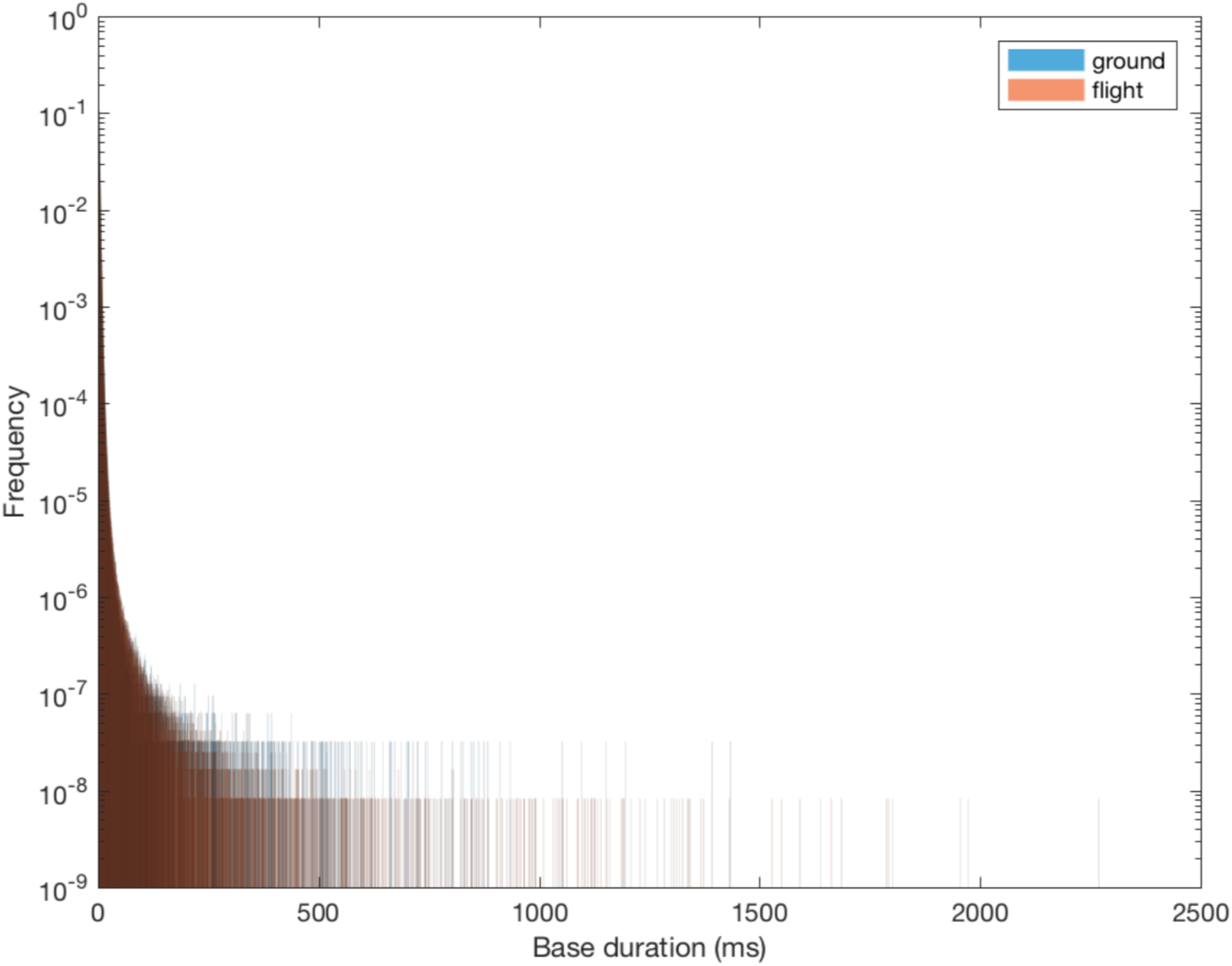
Nanopore translocation time as measured by alignment of ionic current to the genomic reference: full range. Ground (blue), flight (light brown), both (dark brown).

**Supplementary Fig. 7.**
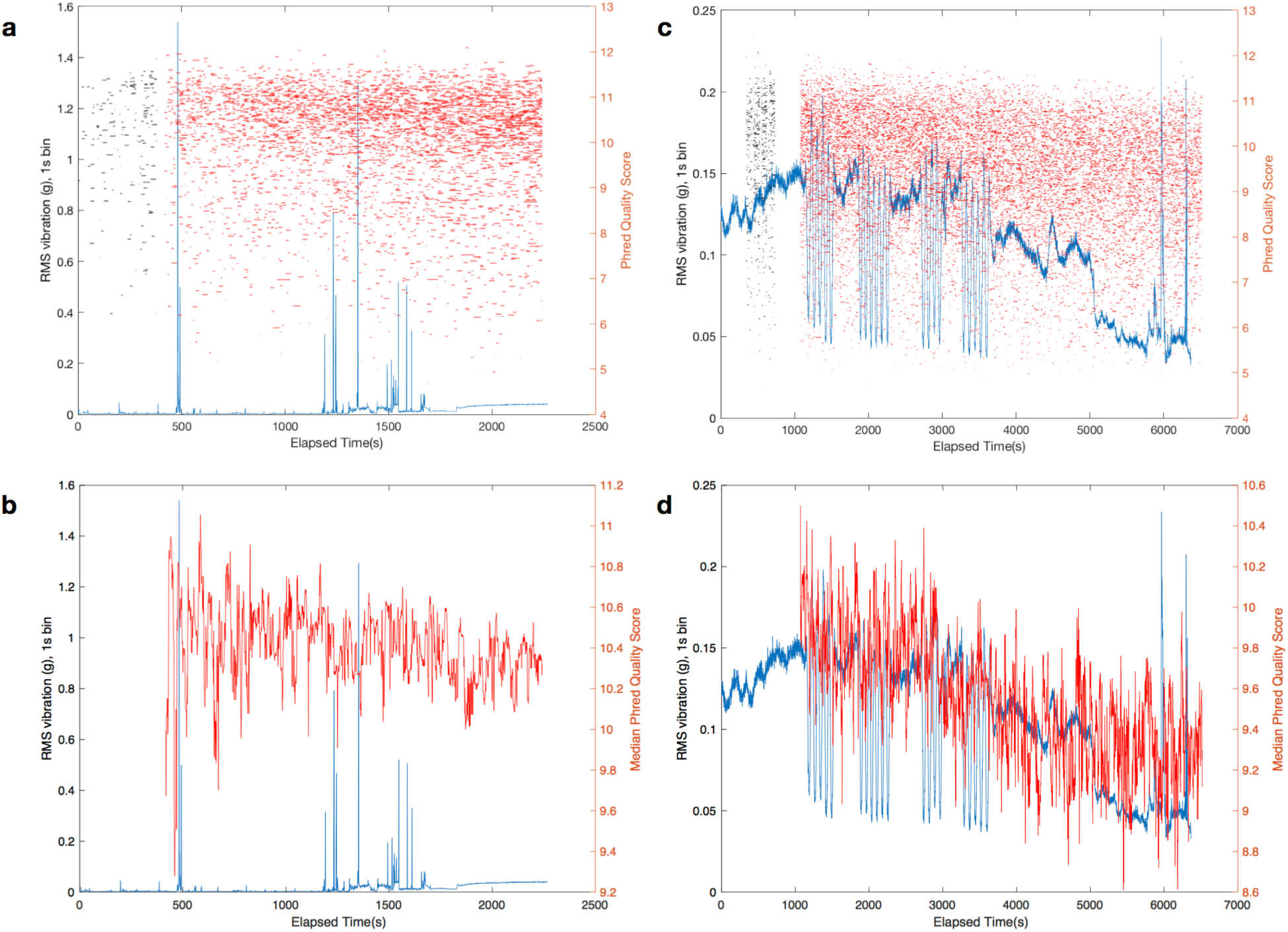
RMS vibration and Sequence Read Quality. **a** Ground RMS vibration (blue) and read quality (mux=grey, run=red). **b** Ground RMS vibration (blue) and median read quality (red). **c** Flight RMS vibration (blue) and read quality (mux=greay, run=red). **d** Flight RMS vibration (blue) and median read quality (red). In panels **a-b**, each horizontal line represents one sequencing read. Mux reads are excluded from panels **b, d**.

**Supplementary Fig. 8.**
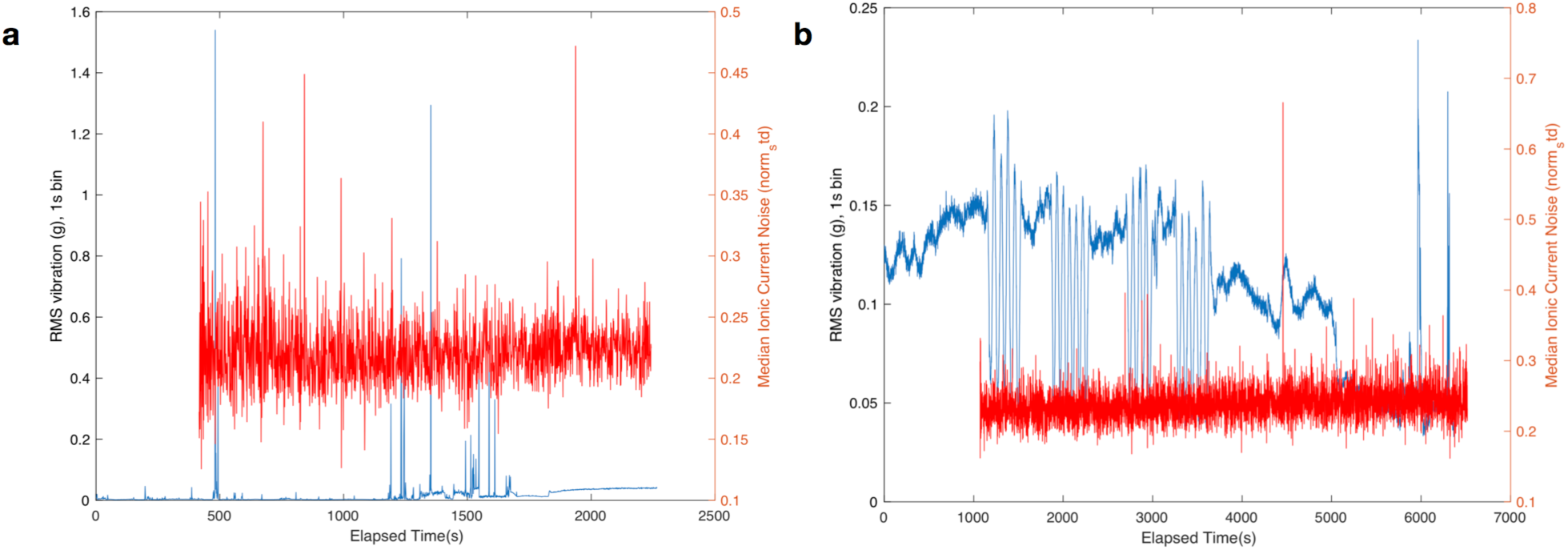
RMS vibration and Ionic Current Noise. **a** Ground RMS vibration (blue) and median ionic current noise (red, 1s bin). **b** Flight RMS vibration (blue) and median ionic current noise (red, 1s bin).

**Supplementary Fig. 9.**
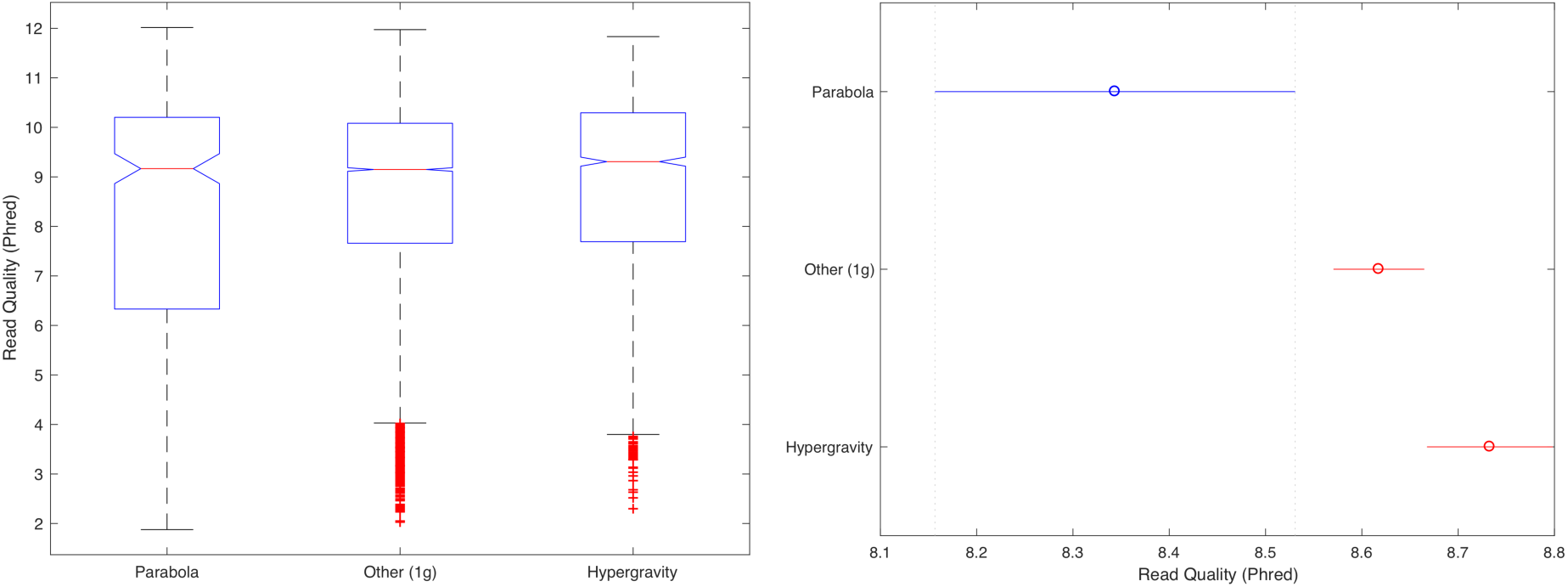
Effect of phase of flight on read quality. Distribution of read quality 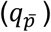 as a function of phase of flight (left) and Tukey Honestly Significant Difference (HSD) test of group means (right). See also: Supplementary Table 5. The “transition” phase of flight is excluded as only 7 reads fell wholly within transition periods.

**Supplementary Fig. 10.**
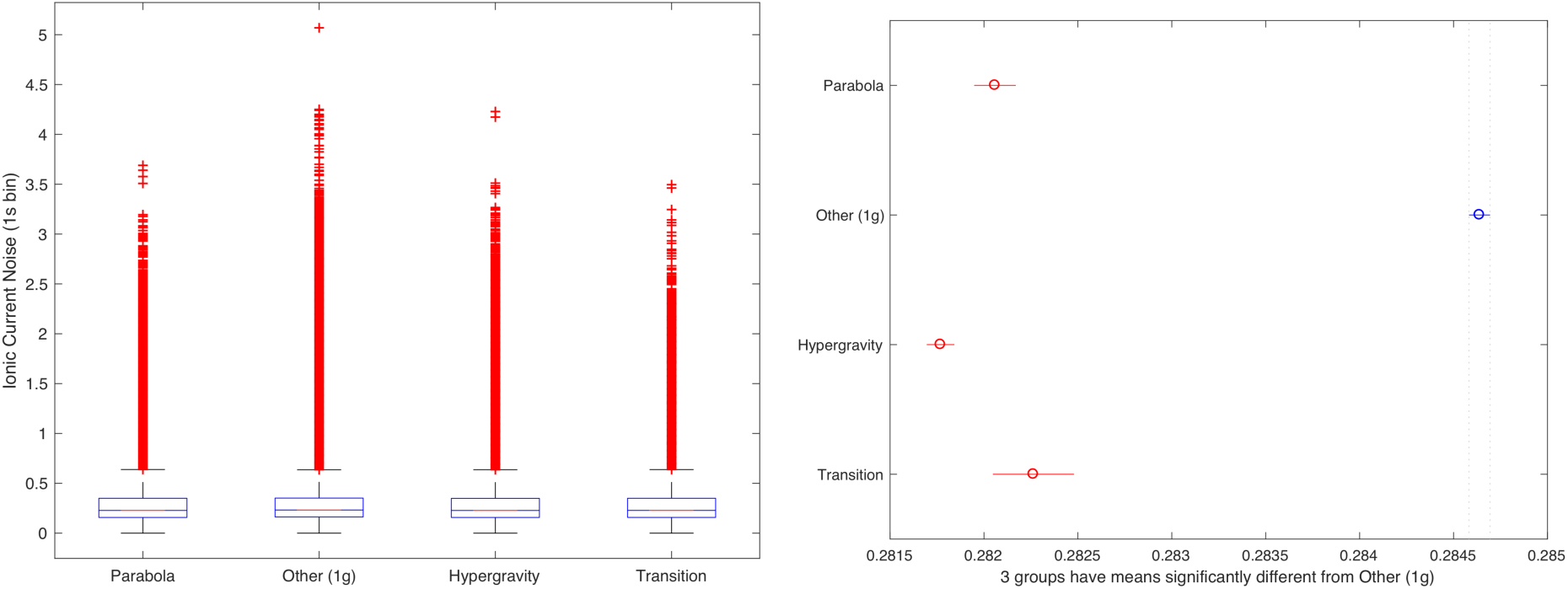
Effect of phase of flight on ionic current noise. Distribution of ionic current noise estimate for each aligned genomic base as a function of phase of flight (left) and Tukey Honestly Significant Difference (HSD) test of group means (right). See also: Supplementary Table 8.

**Supplementary Table 1.**
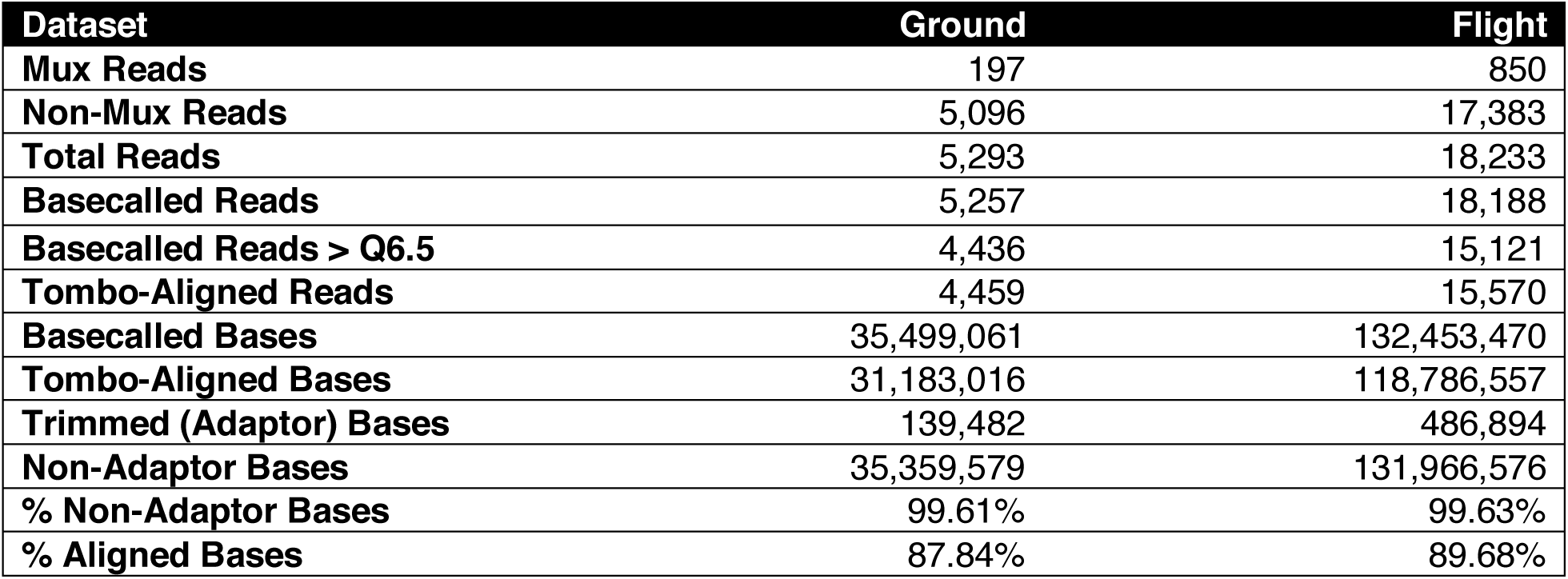
Sequencing Statistics.

**Supplementary Table 2.** Timing Data.

| **Sequencer** | **MinION - Ground** | **MinION - Flight** |
| --- | --- | --- |
| **Mux experiment start time** | 2017-11-17T14:59:03Z | 2017-11-17T18:34:20Z |
| **Run experiment start time** | 2017-11-17T15:06:12Z | 2017-11-17T18:46:32Z |
| **Max read start time in samples (s)** | 7339607 | 21823499 |
| **Corresponding duration (s)** | 4969 | 4981 |
| **Sampling rate (Hz)** | 4000 | 4000 |
| **Sequencing duration (s)** | 1836.144 | 5457.12 |
| **Mux duration (s)** | 429 | 732 |
| **Total sequencing duration (s)** | 2265.144 | 6189.12 |
| **Total sequencing duration (min)** | 37.7524 | 103.152 |
| **Accelerometer** | **SlamStick - Ground** | **SlamStick - Flight** |
| **Start time** | 2017-11-17T14:59:26Z | 2017-11-17T18:28:51Z |
| **Timing Offsets** | **MinION - Ground** | **MinION - Flight** |
| **Mux Offset (s)** | -23 | 329 |
| **Run Offset (s)** | 406 | 1061 |

Timing offsets are added to sequencing time to get accelerometer elapsed time.

**Supplementary Table 3.** Ground Operations: Do time and RMS vibration predict median sequence quality?

| Dataset | Estimate | SE | t-stat | p-value |
| --- | --- | --- | --- | --- |
| Measurements | 1827 |  |  |  |
| Error degrees of freedom | 1825 |  |  |  |
| Regression RMS Error | 0.182 |  |  |  |
| Adjusted R-squared | 0.0602 |  |  |  |
| F-statistic vs. constant model | 118 |  |  | 1.1E-26 |
| Intercept | 10.525 | 0.01157 | 909.63 | 0 |
| Time (s) | -8.7908E-5 | 8.0926E-6 | -10.863 | 1.1E-26 |

Above table is final model from stepwise linear regression. Process from constant model:
pValue for adding Time is 1.1145e-26
pValue for adding Vibration is 0.080905
1. Adding Time, FStat = 117.9991, pValue = 1.114479e-26
pValue for adding Vibration is 0.89155
No candidate terms to remove

**Supplementary Table 4.** Flight Operations: Do time, RMS vibration, and *g*-level predict median sequence quality?

| Dataset | Estimate | SE | t-stat | p-value |
| --- | --- | --- | --- | --- |
| Measurements | 2931 |  |  |  |
| Error degrees of freedom | 2927 |  |  |  |
| Regression RMS Error | 0.200 |  |  |  |
| Adjusted R-squared | 0.275 |  |  |  |
| F-statistic vs. constant model | 371 |  |  | 2.1E-204 |
| Intercept | 9.9413 | 0.028948 | 343.42 | 0 |
| Time | -9.8603E-05 | 1.1189E-05 | -8.8126 | 2.05E-18 |
| g-level | 0.14773 | 0.025729 | 5.7416 | 1.03E-08 |
| Time:g-level | -4.2902E-05 | 9.9399E-06 | -4.3161 | 1.64E-05 |

Above table is final model from stepwise linear regression. Process from constant model:
pValue for adding Time is 2.4094e-197
pValue for adding Vibration is 1.9926e-18
pValue for adding g-level is 0.00017903
1. Adding Time, FStat = 1051.3572, pValue = 2.4093671e-197
pValue for adding Vibration is 1.4909e-06
pValue for adding g-level is 1.2423e-07
2. Adding g-level, FStat = 28.0928, pValue = 1.24226e-07
pValue for adding Vibration is 0.25185

**Supplementary Table 5.** Flight Operations: Does read quality differ between phases of flight?

| **Source** | **SS** | **df** | **MS** | **F** | **Prob>F** |
| --- | --- | --- | --- | --- | --- |
| **Groups** | 55.6 | 2 | 27.8023 | 7.16 | 0.0008 |
| **Error** | 52548.9 | 13531 | 3.8836 |  |  |
| **Total** | 52604.5 | 13533 |  |  |  |
| **Group1** | **Group 2** | **CI (low)** | **HSD** | **CI (high)** | **p-value** |
| **Parabola** | **Other (1g)** | -0.5081 | -0.2739 | -0.0397 | 0.0168 |
| **Parabola** | **Hypergravity** | -0.6415 | -0.3892 | -0.1369 | 0.0009 |
| **Other (1g)** | **Hypergravity** | -0.2277 | -0.1153 | -0.0028 | 0.0431 |

Group means: 8.3437 (parabola), 8.6177 (other), 8.7329 (hypergravity). HSD = Honest Significant Difference. CI is 95% percentile.

**Supplementary Table 6.** Ground Operations: Do time and RMS vibration predict ionic current noise?

| Dataset | Estimate | SE | t-stat | p-value |
| --- | --- | --- | --- | --- |
| Measurements | 1827 |  |  |  |
| Error degrees of freedom | 1823 |  |  |  |
| Regression RMS Error | 0.0267 |  |  |  |
| Adjusted R-squared | 0.00903 |  |  |  |
| F-statistic vs. constant model | 6.55 |  |  | 0.000211 |
| Intercept | 0.21866 | 0.0018111 | 120.74 | 0 |
| Time | 2.1177E-06 | 1.3858E-06 | 1.5281 | 0.12666 |
| Vibration | -0.077314 | 0.024691 | -3.1313 | 0.0017679 |
| Time:Vibration | 4.7315E-05 | 2.3164E-05 | 2.0426 | 0.041235 |

Above table is final model from stepwise linear regression. Process from constant model:
- pValue for adding Time is 0.012543 - pValue for adding Vibration is 0.009305 - 1. Adding Vibration, FStat = 6.7777, pValue = 0.009305 - pValue for adding Time is 0.0033302 - 2. Adding Time, FStat = 8.6399, pValue = 0.0033302 - pValue for adding Time:Vibration is 0.041235 - 3. Adding Time:Vibration, FStat = 4.1722, pValue = 0.041235 - No candidate terms to add - No candidate terms to remove

**Supplementary Table 7.** Flight Operations: Do time, RMS vibration, and g-level predict ionic current noise?

| Dataset | Estimate | SE | t-stat | p-value |
| --- | --- | --- | --- | --- |
| Measurements | 2931 |  |  |  |
| Error degrees of freedom | 2929 |  |  |  |
| Regression RMS Error | 0.0208 |  |  |  |
| Adjusted R-squared | 0.0137 |  |  |  |
| F-statistic vs. constant model | 41.8 |  |  | 1.18E-10 |
| Intercept | 0.22626 | 0.0012148 | 186.26 | 0 |
| Time (s) | 2.9385E-06 | 4.5455E-07 | 6.4647 | 1.18E-10 |

Above table is final model from stepwise linear regression. Process from constant model:
- pValue for adding Time is 1.185e-10 - pValue for adding Vibration is 0.99466 - pValue for adding g-level is 0.28286 - 1. Adding Time, FStat = 41.792, pValue = 1.18499e-10 - pValue for adding Vibration is 0.27136 - pValue for adding g-level is 0.36804 - No candidate terms to remove
Note: extending analysis time to entire flight does not change any of our conclusions.

**Supplementary Table 8.** Flight Operations: Does ionic current noise differ between phases of flight?

| Source | SS | df | MS | F | Prob>F |
| --- | --- | --- | --- | --- | --- |
| **Groups** | 183.6898 | 3 | 61.2299 | 1.6204e+03 | 0 |
| **Error** | 4.2091e+06 | 111393248 | 0.0378 |  |  |
| **Total** | 4.2093e+06 | 111393251 |  |  |  |
| Group1 | Group 2 | CI (low) | HSD | CI (high) | p-value |
| **Parabola** | Other (1g) | -0.00275 | -0.00258 | -0.00241 | 3.77E-09 |
| **Parabola** | Hypergravity | 0.00010 | 0.00029 | 0.00048 | 5.96E-04 |
| **Parabola** | Transition | -0.00052 | -0.00020 | 0.00011 | 0.345 |
| **Other (1g)** | Hypergravity | 0.00275 | 0.00287 | 0.00299 | 3.77E-09 |
| **Other (1g)** | Transition | 0.00210 | 0.00238 | 0.00265 | 3.77E-09 |
| **Hypergravity** | Transition | -0.00079 | -0.00049 | -0.00020 | 8.55E-05 |

Group means: 0.2818 (hypergravity), 0.2821 (parabola), 0.2823 (transition), 0.2846 (other/1g). HSD = Honest Significant Difference. CI is 95% percentile.

**Supplementary Table 9.**
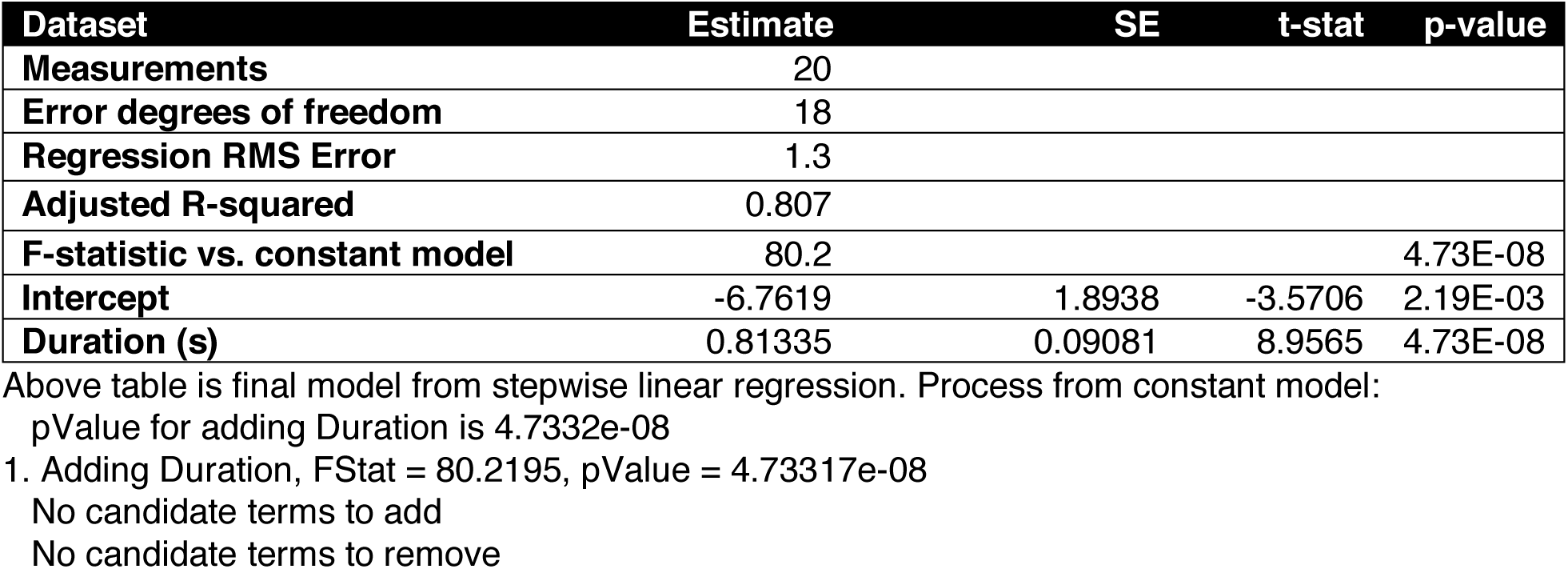
Flight Operations: Does parabola duration predict coverage?

